# Neurons and molecules involved in noxious light sensation in *Caenorhabditis elegans*

**DOI:** 10.1101/2024.12.19.629511

**Authors:** Eva Dunkel, Ichiro Aoki, Alexander Gottschalk

## Abstract

Ultraviolet (UV) light is a danger to unpigmented organisms, inducing photodamage of cells and DNA. The transparent nematode *Caenorhabditis elegans*, despite having no eyes, detects light and exhibits negative phototaxis in order to evade sunlight. UV absorption is detected by the photosensor protein LITE-1, that also responds to reactive oxygen species. We investigated which neurons express LITE-1 and act as noxious photosensors and how they transmit this sensation to the nervous system to evoke escape behavior. We identified the interneuron AVG as a main focus of LITE-1 function in mediating the noxious light evoked escape behavior, with minor roles of the interneuron PVT, the sensory ASK neurons and touch receptor neurons. AVG is activated by blue light, and also its optogenetic stimulation causes escape behavior. Signaling from AVG involves chemical neurotransmission, likely directly to premotor interneurons, and to other cells, by extrasynaptic signaling through the neuropeptide NLP-10. NLP-10 signaling is not required for the acute response, but for maintaining responsiveness to repeated noxious stimuli. The source of NLP-10 in this context is largely AVG, however, also other cells contribute, possibly PVT. This work uncovers entry points of sensory information to the neuronal circuits mediating behavioral responses to noxious UV/blue light.

**Article Summary:** *C. elegans* senses noxious light and induces escape behavior to avoid damage. The photosensor protein LITE-1 mediates this sensation but understanding responsible neural circuits is incomplete. We identified neurons expressing LITE-1 and identify AVG as the main site of action. The neurotransmitter GABA plays a role in acuteness of the response. AVG, and other cells, need to release the neuropeptide NLP-10 to maintain responsiveness to repeated noxious stimuli. These findings help understanding the *C. elegans* photophobic response and will guide future work delineating the precise circuit pathways, as an example of how similar photosensation can evoke protective behavior in invertebrates.

## Introduction

In the animal kingdom, different strategies exist to sense light and to adapt behavior accordingly. The eyes of vertebrates contain millions of photoreceptors, allowing them to sense their surrounding with great precision, which is essential for navigation and decision-making. The signals received by the retina are transmitted to the primary visual cortex via the lateral geniculate nucleus and processed in two streams in the cortex, eventually triggering behavioral output (Goodale and Milner, 1992; Ungerleider, 1982). Insects and arthropods have a comparatively simpler visual system, with thousands of sensory cells and complex downstream neural circuits that allow them to respond quickly, such as adapting their flight behavior to their environment (Land, 1999). Least complex forms of animal vision result from photosensitive neurons and small circuits that can evoke tactic behaviors, for example to avoid damage from UV light. UV-induced DNA damage involves formation of cyclobutane pyrimidine dimers, caused by absorption of UV-B photons (280-315 nm), and is linked to skin cancer development (Cadet and Douki, 2018; Heil et al., 2011; Taylor, 1994). The transparent nematode *Caenorhabditis elegans*, lacking organs for visual sensation, lives in soil or rotting biomass, which protects the animals from sunlight. Should it leave this habitat, negative phototaxis to UV and blue light, regulated by the LITE-1 protein, ensures its return to protected areas (Edwards et al., 2008).

The mechanism of UV light detection by LITE-1 is not yet clarified (Bhatla and Horvitz, 2015; Gong et al., 2016; Xiang et al., 2010). LITE-1 (along with GUR-3, expressed in pharyngeal neurons and affecting light-evoked pharynx inhibition; Bhatla and Horvitz, 2015) is a member of a protein family including insect gustatory / olfactory receptors, which are ion channels (Butterwick et al., 2018; Del Marmol et al., 2021; Saina et al., 2015). Based on these similarities and on Alphafold multimer modeling, LITE-1 was suggested to be a nociceptive light-activated ion channel (Abramson et al., 2024; Evans et al., 2022; Hanson et al., 2023). Moreover, LITE-1 is responsible for locomotion escape behavior, that is either forward acceleration, or a reversal, depending on whether the noxious light hits the posterior or anterior part of the animal, respectively (Edwards et al., 2008). This implies that both anterior and posterior cells mediate the escape response. Animals lacking LITE-1 show a largely decreased escape response. In preceding studies, a general rescue was obtained by expressing LITE-1 under its own promoter in *lite-1(ce314)* mutants (Aoki et al., 2024; Edwards et al., 2008). mRNA of LITE-1 was detected in 15 different cell types in *C. elegans* and appears to be most abundantly expressed in the AVG neuron (Taylor et al., 2021) (**Fig. S1A, Table S1**). This cholinergic interneuron is a pioneer neuron for the right track of the ventral nerve cord (VNC) (Pereira et al., 2015; Riddle et al., 1997). Ablation of AVG in embryos leads to aberrant anatomical distribution of the axons of several ventral cord inter- and motor neuron, causing them to extend on the left side of the VNC (Durbin, 1987; Garriga et al., 1993). The PVT neuron, exhibiting the second highest LITE-1 mRNA levels, is a single interneuron located at the posterior end of the right VNC (Taylor et al., 2021), and is similarly important for normal VNC development (Ren et al., 1999). Removal of PVT in L1 stage animals results in positional changes of axons that were already embedded in the VNC (Aurelio et al., 2002). AVG, PVT and ASK, the latter is a head sensory neuron involved in chemotaxis behavior and ranking third in LITE-1 mRNA levels, were previously implicated as possible sites of action of LITE-1, or as photosensitive neurons (Bargmann and Horvitz, 1991; Edwards et al., 2008; Liu et al., 2010; Setty et al., 2022; Ward et al., 2008). Other LITE-1 expressing cells include touch sensory neurons ALM and PLM.

It is unknown how the nociceptive signal perceived by LITE-1 expressing neurons is relayed to the remainder of the nervous system. Cells that receive innervation by these neurons are candidates for transforming the photoresponse into locomotion behavior (**Fig. S1B**). These include AVA and AVB neurons, downstream of AVG, which are premotor interneurons that instruct reverse and forward locomotion, respectively. Also, the so-called first layer interneurons of the chemosensory navigation circuit, e.g. AIB and AIA neurons, which are innervated by PVT and ASK, relay signals to premotor interneurons (Gray et al., 2005). PVT and the touch receptor neurons also innervate AVK and DVA interneurons that affect navigation (Aoki et al., 2024; Oranth et al., 2018). However, LITE-1 signaling may also mediate a systemic signal, that could alert many cells and tissues of danger by UV light. This idea is supported several findings: 1) LITE-1 activation was shown to suppress the near-paralyzed phenotype of *unc-31* mutants (encoding the Ca^2+^ activator protein for secretion - CAPS), which have reduced synaptic cAMP signaling and largely lack the release of neuropeptides (Edwards et al., 2008; Rupnik et al., 2000; Speese et al., 2007; Steuer Costa et al., 2017; Yu et al., 2021). 2) *lite-1* mutants show abnormal locomotive behavior, such as reduced speed and impaired swimming (Edwards et al., 2008). 3) The absence of LITE-1 facilitates effects of the activation of a photosensitive cholinergic agonist (Damijonaitis et al., 2015). Thus, it appears plausible that LITE-1 may be responsible for the release of signaling molecules that generally affect locomotion. The signaling network of neuropeptides affects the whole organism (Beets et al., 2023; Randi et al., 2023; Ripoll-Sanchez et al., 2023; Smith et al., 2019; Taylor et al., 2021). One of the neuropeptides affecting *C. elegans* locomotion is NLP-10, which is expressed in multiple neurons, including AVG and PVT (Taylor et al., 2021). NLP-10 and its receptor, NPR-35, are required for the efficient response to noxious blue light, particularly to sustain the response upon repeated stimulation (Aoki et al., 2024).

We aimed to identify the site of action for LITE-1 by cell-specific rescue. Our results suggest that AVG is the main site of action for the LITE-1 induced escape behavior in response to blue light exposure. The escape behavior was restored to wild type level when rescuing LITE-1 in AVG, while suppressing neurotransmitter release from AVG resulted in impaired escape behavior. AVG Ca^2+^ levels are increased upon blue light illumination, thus underlining the intrinsic reactivity of this neuron, whereas this reaction was abolished in *lite-1* mutants. We further identified AVG as a source of NLP-10 in this context, as specific expression of NLP-10 in AVG partially repressed habituation to repetitive blue light stimulation. However, effective suppression of habituation required additional neurons to express LITE-1, i.e., ASK and PVT, the latter of which also expresses NLP-10. LITE-1 expressing neurons thus affect locomotion likely via synaptic networks and retain nociceptive sensitivity via the NLP-10 neuropeptide.

## Results

### The interneuron AVG is the main site of action of LITE-1

To identify possible sites of action for LITE-1, we expressed the protein in *lite-1(ce314)* mutants under different promoters that are specific or even exclusive for neurons that natively express the LITE-1 gene (**Fig. S2**). These cells include, based on single-cell RNAseq data and by abundance of the *lite-1* mRNA (Taylor et al., 2021), the neurons AVG, PVT and ASK, as well as the touch receptor neurons (TRNs) ALM and PLM (**Fig. S1A, Table S1**). Lower abundance was reported for ASG, PHA, PHB, RIF, RMD, and a number of pharyngeal neurons. We restricted our analysis to the cells with most abundant *lite-1* expression, AVG, PVT, ASK, and the TRNs. The following promoters were cloned and assessed for their expression by transcriptional GFP fusions (**Fig. S2**): The second intron of the *inx-18* gene (*inx-18-int2*) (Oren-Suissa et al., 2016) instructed expression of GFP in a single neuron with the cell body in the retrovesicular ganglion, and a process reaching all along the VNC toward the tail (**Fig. S2A**). The expression matched the reported morphology of AVG (White et al., 1986). A part of the third intron of the *lim-6* gene (*lim-6-int3s)* (Chien et al., 2017) drove expression in a neuron with a cell body in the pre-anal ganglion, and extended a process through the VNC and around the nerve ring, where it formed several varicosities (**Fig. S2B**). This morphology is in line with details reported for the PVT neuron (White et al., 1986; Witvliet et al., 2021), and localization relative to a marker for the AIY neuron corresponded with this interpretation. For the ASKL/R neurons, we tested several promoters based on scRNAseq data (Taylor et al., 2021). We settled for the *ZC190*.*6* promoter, which expressed in two neurons with cell bodies in the lateral nerve ring ganglia, and processes to the amphid sensory organs (**Fig. S2C**). Also, these cells stained positive for the lipophilic dye DiI, which can enter the sensory endings of these neurons. Yet, two additional, unknown neurons, one in the tail and one about halfway between the pharyngeal terminal bulb and the vulva (possibly, ALM or AVM neurons, based on position and morphology) were expressing this promoter. For the TRNs, we used the well-established *mec-4* promoter (Chalfie and Sulston, 1981).

We then assessed transgenic animals expressing LITE-1 cDNA from these promoters for their potential to rescue the blue-light evoked escape response. As controls, we used wild type animals (or rather, the *juSi164* background, used for transgene integration into the genome; *juSi164* expresses the miniature single oxygen generator (miniSOG) fused to histone HIS-72; Noma and Jin, 2018), as well as *juSi164; lite-1(ce314)* mutants. During the blue light stimulation (1 mW/mm^2^), *lite-1* animals showed an increase in speed (1.2 fold) that was significantly lower than that of wild type animals, which increased their speed about 4 fold (**Fig. 1A-C** note, data were normalized to the respective mean during the 270-299 s just preceding the light stimulus). This difference was not only detectable in statistical analysis but also upon visual examination of the animals’ behavior (**Videos S1, S2**). *lite-1* mutants expressing LITE-1 in AVG showed a clear increase in speed, up to 3.9-fold, i.e. a significant, complete rescue of the light-evoked escape behavior. For animals expressing LITE-1 in the PVT neuron, there was only a minor initial speed increase upon noxious blue light. Thus, PVT contributes only to a minor extent to the light-evoked escape behavior. Strains expressing LITE-1 in AVG presented a slightly delayed onset of the reaction, however, the maximum speed during the illumination period was not reached significantly later than in wild type controls (**Fig. 1D, E**). The time to reach the maximum speed during the illumination period for animals carrying PVT::LITE-1 likewise did not differ from wild type or AVG::LITE-1 animals. However, PVT::LITE-1 animals exhibited a pronounced, clearly delayed offset response following the end of the light stimulus by about 40 sec (**Fig. S3**). Analysis of the reversal rate of animals before, during and after the illumination period in 10 s bins showed increased reversal behavior of *lite-1* animals, indicating some residual light responsiveness, possibly through GUR-3 (Bhatla and Horvitz, 2015), but otherwise did not reveal clear differences between rescued animals and wild type (**Fig. S4A-C**).

**Figure 1:**
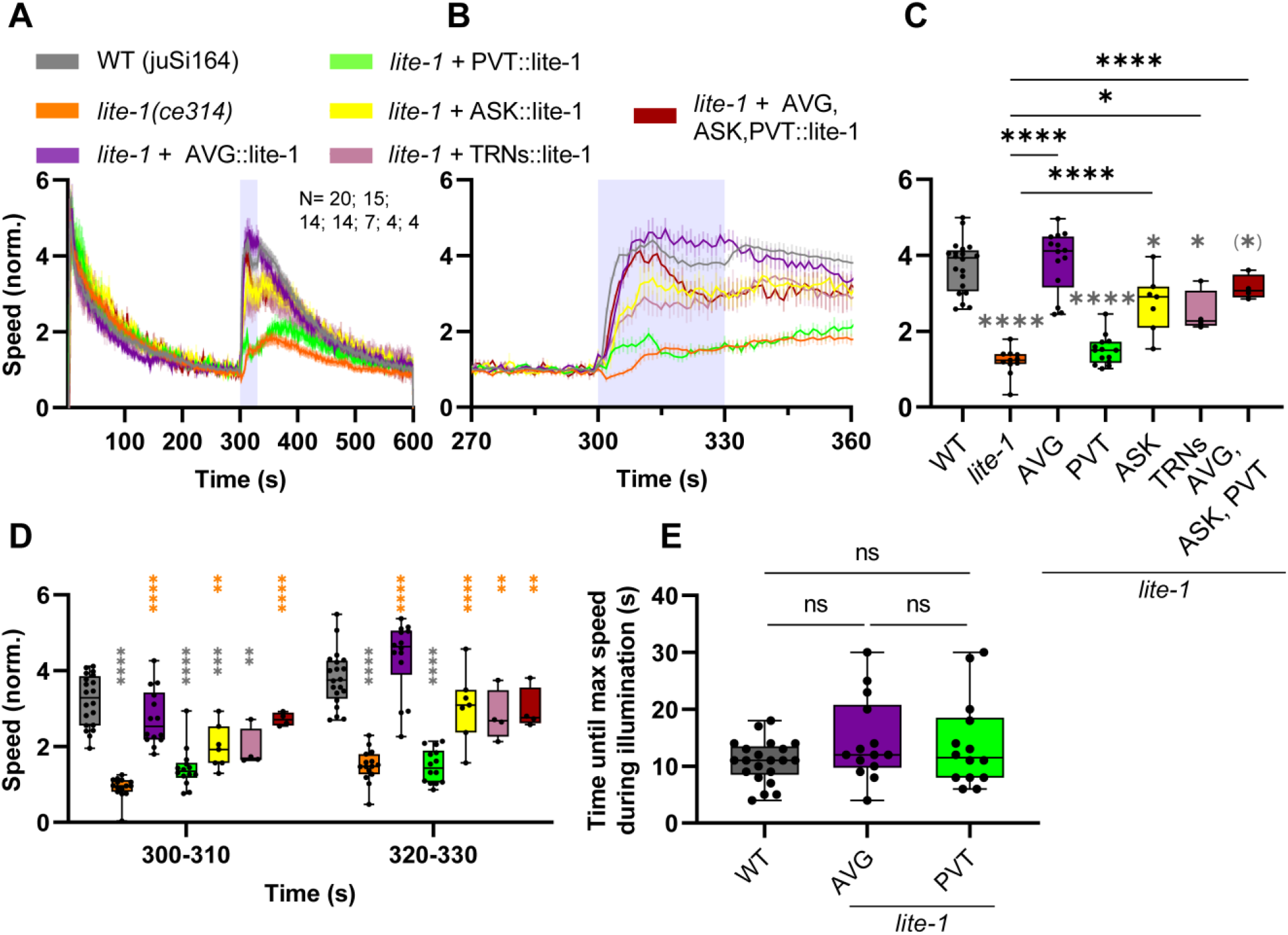
LITE-1 is required mainly in AVG, and in ASK and TRNs as secondary site of action for noxious light-evoked escape behavior. (**A**) Mean crawling speed (± SEM) of animals before (0-299 s), during (300-330 s) and after (331-600 s) blue light illumination. Tested strains were wild type, *lite-1(ce314)* and *lite-1* mutants expressing LITE-1 solely in neurons AVG, PVT, ASK, and in touch receptor neurons (TRNs), or simultaneously in AVG, PVT and ASK. Speed was normalized to the mean crawling speed from 270 to 299 s. Illumination is indicated by the blue shade. N=20, 15, 14, 14, 7, 4, and 4 experiments, with n=20-32 animals each, respectively. (**B**) Close-up of 270-360 s, including the illumination period. (**C**) Comparison of speed among the indicated strains during the illumination period. Median with 25/75 quartiles, whiskers indicate minimum and maximum values. One-way ANOVA with Tukey test (p < 0.05 = *; p < 0.01 = ****). Grey asterisks indicate significant difference to wild type, others as indicated. Note, in a comparison of the time windows of 305-320 s, the triple rescue strain was not different to wild type. (**D**) Comparison of speeds between strains during the first and last ten seconds of illumination. Median with 25/75 quartiles, whiskers indicate minimum and maximum values. Two-way ANOVA with Tukey test. Grey and orange asterisks: Significant differences to wild type and *lite-1*, respectively. (p < 0.05 = *; p < 0.01 = **; p < 0.001 = **; p < 0.0001 = ****). (**E**) Time until maximum speed is reached during illumination period. Median with 25/75 quartiles, whiskers indicate minimum and maximum values. One-way ANOVA with Tukey test (ns = not significant).

Next, we tested the ASK neurons and the TRNs, including PLM and ALM, as possible sites of action for LITE-1. Both rescue strains showed a speed increase upon blue light stimulation that was significant, yet not as pronounced as for wild type animals (**Fig. 1A-D**). Last, we tested a strain expressing all three rescue constructs, i.e. for AVG, PVT and ASK. The combined expression of LITE-1 in neurons AVG, PVT and ASK led to escape behavior that was almost identical in its amplitude and speed development as in wild type animals, resulting in a clear rescue of the *lite-1* phenotype (**Fig. 1A-D**). However, the AVG single neuron rescue was somewhat more similar to wild type (note, the triple rescue was not a chromosomally integrated transgene, in contrast to the single neuron rescue; depending on the time window of analysis, i.e., 305-320 s, the triple rescue was not significantly different to wild type). Given the extent of rescue, AVG seems to be a main focus of activity of LITE-1.

### AVG responds to blue light, depending on the presence of LITE-1

To verify that endogenous LITE-1 function can activate AVG, we examined the effect of blue light on AVG Ca^2+^ levels. The red fluorescent calcium indicator RCaMP, which can be combined with blue light stimulation (Akerboom et al., 2013), showed a significant increase in signal intensity when the animals were illuminated with blue light (**Fig. 2A-C**), demonstrating blue light causes an activation of AVG, leading to a rise of the Ca^2+^ concentration. In *lite-1* mutants, the RCaMP signal increase was absent (**Fig. 2B, C**). Thus, AVG gets activated by blue light, and this effect depends on LITE-1. A negative control expressing a calcium-insensitive fluorescent protein did not change in its signal intensity (**Fig. S5**).

**Figure 2:**
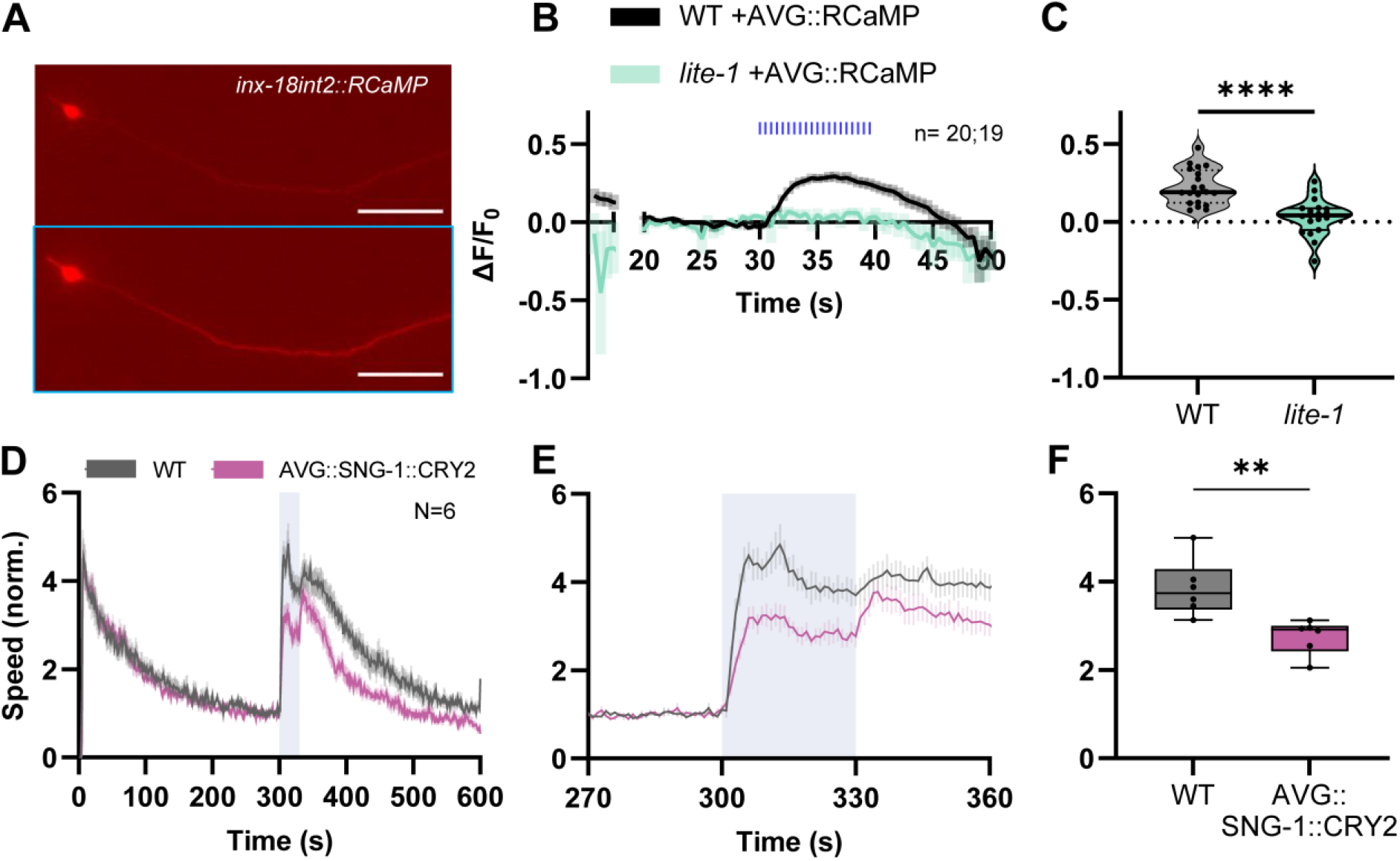
The AVG neuron is activated by blue light and releases a signal by chemical neurotransmission. (**A**) Exemplary images of RCaMP signal in AVG (cell body and axon, anterior is left) before (top) and right after (bottom, blue box) blue light application. Scale bars: 50 µm. (**B**) Mean (±SEM) change of fluorescence intensity of RCaMP in AVG before (20-29 s), during (30-39 s) and after (40-50 s) application of blue light pulses (indicated by blue bars). n=20 animals for wild type and n=19 for *lite-1*. F_0_ was calculated during seconds 20-29. (**C**) Mean RCaMP signal intensity during blue light application of individual animals. Median (thick line) and 25/75 quartiles (dotted or thin lines). Unpaired t-test (p < 0.0001 = ****). (**D**) Mean crawling speed (± SEM) of animals before (0-299 s), during (300-330 s) and after (331-600 s) blue light illumination. Tested strains were wild type and animals expressing optoSynC (optogenetic tool for immobilization of synaptic vesicles) in AVG. Speed was normalized to the mean crawling speed from 270 to 299 s. Illumination indicated by blue shade. N=6 experiments with 27-38 animals each. (**E**) Close-up of locomotion speed 30 s before, during and after illumination. (**F**) Comparison of crawling speed during illumination. Median with 25/75 quartiles, whiskers indicate minimum and maximum values. Unpaired, two-tailed t-test (p = 0.0052).

To further analyze how AVG propagates the light-induced signal to downstream cells affecting locomotion, we targeted chemical synaptic transmission. We inhibited the mobility of synaptic vesicles and thus their recruitment for transmitter release at the active zone plasma membrane, specifically in AVG, using the <u>opto</u>genetic <u>syn</u>aptic vesicle <u>c</u>lustering tool OptoSynC, consisting of the SV protein SNG-1 and CRY2olig, a light-dependent oligomerizing variant of cryptochrome 2 (Vettkotter et al., 2022). We previously showed that pan-neuronal expression of this tool suppressed the reaction to strong blue light to a level similar as in *lite-1* mutant background (Vettkotter et al., 2022). By expressing optoSynC specifically in AVG and activating it by blue light, we could reduce neurotransmitter release from AVG. This led to a significantly decreased escape behavior, yet did not abolish it (**Fig. 2D-F**). Potentially, optoSynC may have targeted transmission by neuropeptides from dense core vesicles, provided SNG-1 is part of these vesicles, too.

### Depolarization of AVG leads to increased locomotion speed, which is affected in absence of GABA

LITE-1 was recently suggested to be a light-activated ion channel (Hanson et al., 2023). This implies that depolarization of AVG should cause the same behaviors as activation of LITE-1. At the same time, it would demonstrate that AVG-specific expression of the LITE-1 protein is inducing cellular signaling based on depolarization. We thus used the AVG promoter to drive expression of Chrimson, a red shifted channelrhodopsin (ChR) variant (Klapoetke et al., 2014; Oda et al., 2018), specifically in AVG (**Fig. S6**). When animals expressing AVG::Chrimson were fed the ChR chromophore all-*trans* retinal (ATR) and illuminated with red light (620 nm, 0.3 mW/mm^2^), we observed a clear increase in locomotion speed (**Fig. 3A, B; Video S3**). This effect was not observed in animals that were not treated with ATR beforehand, ruling out any behavioral effect caused by the red light illumination on the animals (**Fig. 3C**).

**Figure 3:**
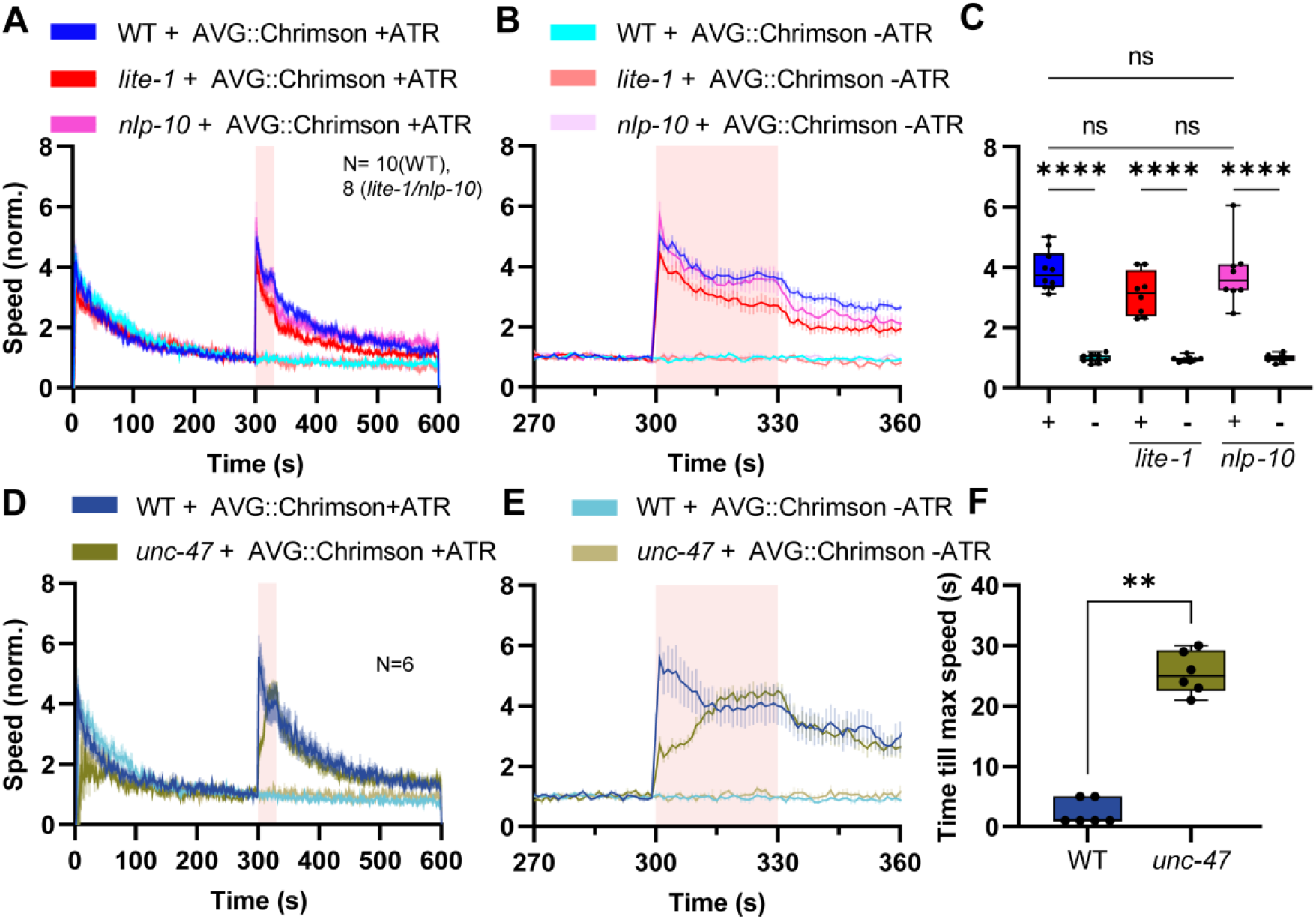
Optogenetic stimulation of AVG triggers escape behavior. (**A**) Mean crawling speed (± SEM) of animals expressing Chrimson in AVG in wild type (N=10), *lite-1(ce314)* (N=8) and *nlp-10(zx29)* (N=8 with 19-35 animals for each strain) were analyzed before, during and after red light illumination (red shade). Crawling speeds were normalized to the respective mean crawling speeds from 270 to 299 s. Animals without exposure to all-*trans* retinal (ATR) served as a negative controls. (**B**) Close-up of locomotion speed 30 s before, during and after illumination. (**C**) Statistical analysis of data in A, B), during red light illumination. Median with 25/75 quartiles, whiskers indicate minimum and maximum values. Two-way ANOVA with Tukey post-hoc analysis (ns= not significant; p < 0.0001 = ****). (**D, E**) As in A, B, but for wild type or *unc-47(e307)* animals expressing AVG::Chrimson that were either exposed to ATR or not. N=6 with 26-31 animals each. (**F**) Time until animals reached their maximum speed during the illumination. Median with 25/75 quartiles, whiskers indicate minimum and maximum values. Mann-Whitney test (p = 0.0022).

We compared the effect of AVG depolarization in *lite-1* and *nlp-10* mutants. NLP-10 was shown to contribute to the effects of blue light stimulation of animals, particularly to keep up a strong effect during repeated stimulation. For both mutant backgrounds, AVG::Chrimson activation evoked a robust and immediate reaction of animals (**Fig. 3A-C**). Even though *lite-1* mutants showed a slightly weaker reaction upon AVG activation, this difference was not statistically significant, and the responses were observed only in animals treated with ATR. This verifies that AVG depolarization evokes similar behaviors like activation of LITE-1 in AVG, and that AVG is a major focus of function of LITE-1, in line with the hypothesis that LITE-1 is a depolarizing ion channel. The nature of the transmitter that AVG uses to evoke the acute signal is as yet unclear.

AVG was reported to be cholinergic, however, a recent study revealed that AVG also expresses the vesicular GABA transporter (vGAT) UNC-47 (Wang et al., 2024). Thus, to assess if the (mostly) inhibitory neurotransmitter GABA might contribute to the AVG-evoked escape behavior, we crossed AVG::Chrimson into *unc-47(e307)* mutants. Upon red light illumination and therefore Chrimson activation, an increase in speed was still detectable (**Fig. 3D-F**). Nevertheless, the behavior of *unc-47* animals differed from wild type animals: The immediate reaction to the stimulation was weaker and animals slowly increased their speed level, reaching the peak much later than wild type animals (**Fig. S7; Video S4**). These results suggest that GABA might facilitate the immediate response upon AVG activation, though it is not essential for it. Since our readout is based on locomotion, and GABA signaling is crucial for proper function of the neuromuscular junction (Bamber et al., 1999), we tested if *unc-47* mutants are generally affected for LITE-1 induced behavioral responses. *unc-47* animals had reduced basal locomotion speed (**Fig. S8D, E**), so we normalized the data to the time before the light stimulus was presented (**Fig. S8A, B**). While wild type animals showed the previously observed rapid and robust (ca. 4-fold) speed increase, this response was attenuated for *unc-47* mutants, which increased their speed only about 2-fold, though they increased their speed significantly more than *lite-1* mutants (**Fig. S8A-C**). The *unc-47* mutant animals showed a much larger offset speed increase after the end of the blue light stimulus, for unknown reasons (**Fig. S8A, B**). In sum, GABA signaling affects the acuteness of LITE-1 evoked escape responses, either directly or indirectly.

### AVG-derived NLP-10 rescues the behavioral habituation effect to repeated blue light stimulation

As we showed recently, the NLP-10 neuropeptide suppresses the habituation to repetitive strong blue light stimulation (Aoki et al., 2024). Since AVG (and PVT) also express NLP-10 (**Table S1, Fig. S1A**), it is possible that NLP-10, released from AVG upon noxious blue light stimulation, contributes to the sustained responsiveness to the noxious stimulus, as long as it persists. To examine the contribution of NLP-10, released from AVG, we expressed mScarlet, fused to the prepropeptide of NLP-10, in AVG. We verified the expression pattern and found that, as expected, mScarlet signal was visible in the AVG cell body. Signal was also observed in coelomocytes, which are scavenger cells that endocytose and filter the body fluid (Sieburth et al., 2007), thus demonstrating that NLP-10 is released from AVG (**Fig. 4A**). We then subjected animals to repeated blue light exposures and analyzed their locomotion behavior (**Fig. 4B-F**). During the first illumination, *nlp-10(zx29)* mutants and *nlp-10* mutants expressing NLP-10 only in AVG showed responses similar to wild type animals (**Fig. 4B-D**), however, the *nlp-10* mutants (and, to some extent, the AVG-specific NLP-10 rescue animals) showed a slower onset of the response. Thus, NLP-10 may be required for the acuteness of the blue light escape behavior, though the source of NLP-10 may not be exclusively AVG. As observed before (Aoki et al., 2024), the extent of speed increase became smaller for each consecutive stimulus in *nlp-10* mutants (**Fig. 4B-F**), while it was maintained at the high level of wild type animals also in the *nlp-10* animals with NLP-10 rescue in AVG. To more directly analyze this, we subtracted the mean speed of the 4^th^ stimulus from that of the 1^st^ stimulus (**Fig. 4F**), showing that the AVG::NLP-10 animals were significantly better than *nlp-10* mutants, and not different from wild type. Overall, AVG-derived NLP-10 led to a rescue of enhanced habituation to noxious blue light evoked escape behavior in *nlp-10* mutants. Given the time course of the speed increase during illumination, AVG is likely not the only site of action of NLP-10 regarding this phenotype (Aoki et al., 2024).

**Figure 4:**
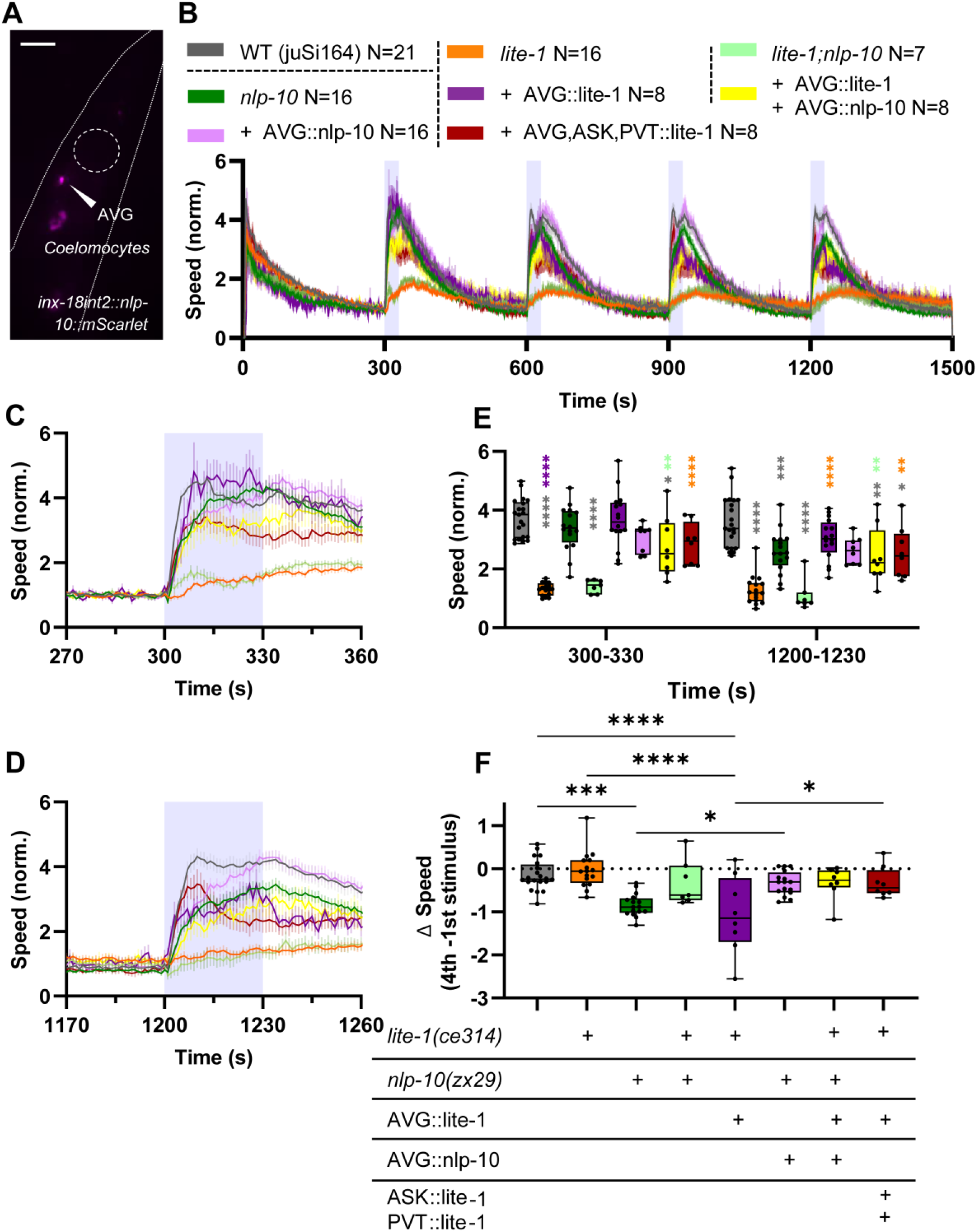
Expression of NLP-10 in AVG rescues habituation to repetitive blue light stimulation. (**A**) Representative image of NLP-10::mScarlet fluorescence, expressed specifically in AVG. White arrowhead indicates AVG soma, additional signal was seen in coelomocytes. Scale bar indicates 50 µm. (**B**) Mean crawling speed (± SEM) of the indicated strains during repetitive blue light stimulation at 300-330, 600-630, 900-930 and 1200-1230 s, as indicated by blue shades. Crawling speed was normalized to the mean crawling speed from 270 to 299 s. N experiments per strain, as indicated, with 23-41 animals each. (**C, D**) Close-ups of locomotion speed 30 s before, during and after the first and the fourth illumination period, respectively. (**E**) Comparison of crawling speed during the first (300-330 s) and fourth illumination (1200-1230 s). Median with 25/75 quartiles, whiskers indicate minimum and maximum values. Two-way ANOVA with Tukey post-hoc analysis was applied (p < 0.05 = *; p < 0.01 = **; p < 0.001 = ***; p < 0.0001 = ****). Color of asterisks indicates significant differences to the respective strain. (**F**) Differences of mean speeds of fourth illumination, deduced by speed levels at first illumination. Median with 25/75 quartiles, whiskers indicate minimum and maximum values. One-way ANOVA with Tukey test (p < 0.05 = *; p < 0.01 = **; p < 0.001 = ***; p < 0.0001 = ****).

We further analyzed this effect in *lite-1; nlp-10* double mutants, as well as in cell specific rescue strains for both genes. As expected, the double mutants showed only a minimal speed increase upon the first light stimulation, just as *lite-1* mutants (**Fig. 4B-F**). The reduced responsiveness of the *lite-1* or the *lite-1; nlp-10* double mutants for the first stimulus was partially rescued by expressing LITE-1 in AVG. However, expressing both LITE-1 and NLP-10 in the double mutant reduced the overall response. Animals expressing LITE-1 and NLP-10 in AVG in the respective double mutant showed only a partial rescue of the response. However, they appeared to maintain their sensitivity to repetitive stimulation, as the habituation levels were similar to wild type. Comparing the habituation (4^th^ minus 1^st^ stimulus; **Fig. 4F**) showed that while rescuing LITE-1 in AVG in *lite-1* mutants led to a robust rescue of the behavior, the speed difference between the fourth and first stimulation was significantly less than wild type, a similar effect as in *nlp-10* mutants. Therefore, we asked if other cells expressing LITE-1 and/or NLP-10 may be required for maintaining full responsiveness such as in wild type. We thus rescued LITE-1 in AVG, PVT and ASK simultaneously. This maintained the nociceptive responsiveness upon the 4^th^ stimulus significantly better than *lite-1* mutants expressing LITE-1 in AVG only. To probe if NLP-10 release is also required to maintain responsiveness to AVG photostimulation using AVG::Chrimson, we compared the behavioral responses in wild type and in *nlp-10* mutants (**Fig. S9**). AVG stimulation caused robust escape behavior following each of four stimuli. In *nlp-10* mutants, this was slightly, but significantly, affected. In sum, blue light evoked nociceptive behavior largely involves the stimulation of AVG, and to some extent also ASK and/or PVT. Furthermore, AVG is a source of NLP-10, enabling to maintain nociceptive responsiveness. However, other cells that release NLP-10, likely including PVT, are required to fully maintain the blue light (AVG) -evoked locomotion speed increase.

## Discussion

Here, we analyzed which neurons contribute to the UV-blue light avoidance of *C. elegans*. The neuron AVG, which expresses highest levels of LITE-1 mRNA, is also the most important neuron driving the escape behavior. Other cells, particularly ASK and touch receptor neurons contribute to this response. PVT, the neuron expressing the second highest amount of *lite-1* mRNA is not required for the acute response, but may contribute to the overall maintenance of the nociceptive responsiveness upon repeated exposure. This aspect of LITE-1 induced signaling requires the neuropeptide NLP-10, for which a major source in this context is again AVG. Which signal is the driver for the acute locomotion speed increase, however, remains unclear. It is likely to be sought in the neuronal circuits downstream of AVG, specifically the premotor interneurons for forward locomotion. However, AVG also forms synapses with backward premotor interneurons. It remains to be identified how the two signaling aspects may be coordinated. The fact that AVG appears to be both cholinergic and GABAergic could imply specific, opposing signal output to the respective interneuron type, promoting net forward acceleration. However, at stage, this is speculation.

Intense short-wave light causes damage in living organisms, which can even be fatal. Recent studies describe LITE-1 as an ion channel, which is light activated, and acts as a receptor for photons and H_2_O_2_-coincidence via PRDX-2 (Hanson et al., 2023; Quintin et al., 2022). Another member of the gustatory receptor family of odorant receptors of is GUR-3 (Bhatla and Horvitz, 2015). The site of action of these proteins is partially known. Spatial resolution of the photosensation does not occur by a light-sensitive organ, but by the location of the neurons and/or their processes, potentially in varicosities, expressing LITE-1 and GUR-3 along the body. I.e. GUR-3 is restricted to neurons of the pharyngeal nervous system and mediates a stop of pharyngeal pumping, along with LITE-1 (Bhatla and Horvitz, 2015). As LITE-1 is only expressed in a handful of neurons, our goal was to identify its site of action in the context of the blue light induced escape behavior.

Previous work implicated AVG, PVT and ASK in the context of (*lite-1* mediated) negative phototaxis based on expression of a reporter construct and on cell ablation experiments (Edwards et al., 2008; Liu et al., 2010; Ward et al., 2008). When noxious light is applied on the posterior part of the animal, AVG might evoke escape behavior, whereas in the anterior end of the animal, PVT (but not AVG) extends into the nerve ring, suggesting that PVT may mediate LITE-1 mediated reversal behavior. ASK ablation (alongside other ciliated neurons such as ASJ, AWB and ASH that extend amphid sensory endings to the nose tip) resulted in a reduced reversal responses, whereas LITE-1 rescue in ASK in *lite-1* mutants lifted the intensity of photophobic behavior to about 50 % of wild type animals (Liu et al., 2010). In our hands, the expression of LITE-1 exclusively in AVG rescued most of the missing photophobic acceleration response of *lite-1* mutants.

The cholinergic neuron AVG also expresses the vGAT UNC-47, which is involved in GABA and possibly glycine transport (Eastman et al., 1999; Gendrel et al., 2016; McIntire et al., 1993). Photostimulation via Chrimson in AVG resulted in an immediate speed increase in wild type, as well as in *lite-1* and *nlp-10* mutants, however *unc-47* mutants responded with an initially reduced speed increase. Thus, GABA (or glycine) might play a role in the translation of AVG activation to locomotion output. The delay in time until the maximum speed was reached is not caused by the generally uncoordinated crawling behavior of these mutants, since exposing *unc-47* animals to blue light resulted in an immediate onset of the response as in wild type, though with reduced amplitude. Nevertheless, to rigorously demonstrate an involvement of GABA in AVG signaling will require analyzing animals with a cell-specific knock down of *unc-47* mRNA or a cell-specific rescue in an *unc-47* mutant background. Silencing AVG chemical transmission by clustering of synaptic vesicles reduced the reactivity of the animals to noxious light. Previously, it was suggested that LITE-1 activation somehow stimulates cholinergic neurons (Edwards et al., 2008), however, it was unclear whether this was a direct or indirect effect, due to release of signaling molecules that act at the organismic level to activate cholinergic signaling. LITE-1 was found to suppress the activity of a blue-light (470 nm) photoswitchable version of acetylcholine, AzoCholine, when animals are pre-exposed to UV light (350 nm), possibly due to a systemic signal released in response to LITE-1 activation (Damijonaitis et al., 2015). Interestingly, *unc-31* mutants that have an impaired release of neuropeptides and are almost paralyzed, display a normal reaction to blue-violet light (Edwards et al., 2008), thus the signal may not be a neuropeptide. Yet, given the AVG::optoSynC results, it should be release by vesicular exocytosis.

In sum, nociceptive signaling downstream from AVG might mostly involve cholinergic signaling, as well as NLP-10 neuropeptides to sustain the response. NLP-10 might have long-range effects (Ludwig and Leng, 2006; van den Pol, 2012). Our previous study suggests that NLP-10 is crucial to inhibit habituation to noxious blue light stimulation, which could be rescued by the expression of the neuropeptide in different neurons, including AVK (Aoki et al., 2024) that is innervated by PVT via chemical and electrical synapses. Also expression of NLP-10 in AVG rescued the compromised speed increase noticed in *nlp-10* mutants, indicating that AVG is a site of action for NLP-10. Interestingly, a similar effect as seen in *nlp-10* mutants was observed in animals expressing LITE-1 only in AVG. However, when LITE-1 was expressed in AVG, ASK and PVT, animals maintained their responsiveness. This implies that while AVG is the main contributor to the initial response to noxious light avoidance, PVT and/or ASK are crucial for keeping up sensitivity.

## Materials and Methods

### Molecular Biology

The following constructs were generated and/or used in this work:

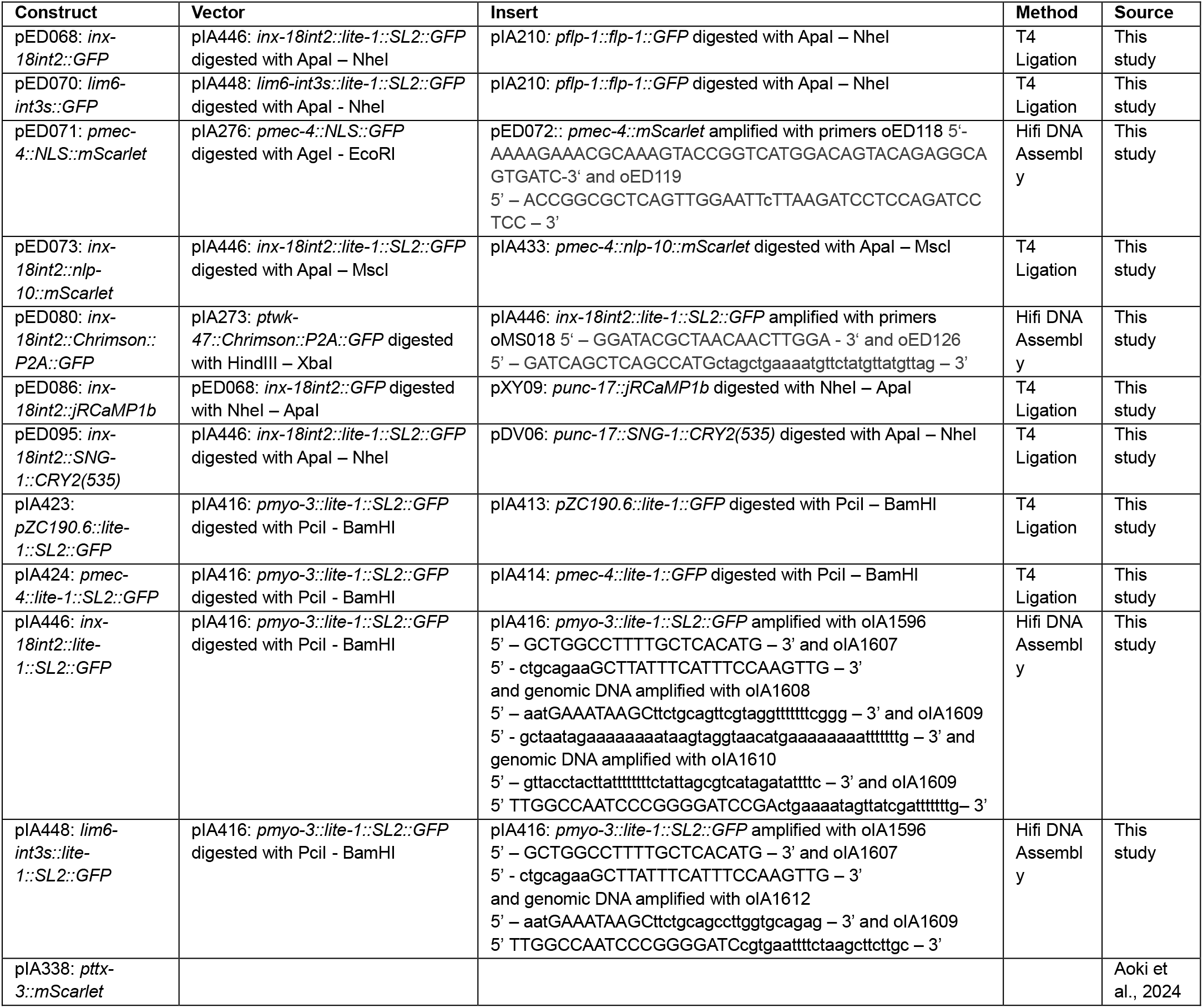

### *C. elegans* strains

*C. elegans* were cultivated on nematode growth medium (NGM) plates seeded with *E. coli* OP-50 bacteria (Brenner, 1974). Unless described otherwise, histone-miniSOG animals (CZ20310 strain carrying *juSi164[pmex-5::his-72::miniSOG + Cbr-unc-119(+)] unc-119(ed3))* were labeled as WT. Strains used in this study are listed below.

For the generation of transgenic lines, plasmid DNA in varying concentrations was injected into the gonads of adult hermaphrodites (Mello et al., 1991). Integration of plasmid DNA into the genome was achieved via activation of histone-miniSOG with blue light illumination 6 hours after injection or after establishing an extrachromosomal strain (Noma et al., 2018). Outcrossed *zxIs270* animals were shown to have identical behavior. An Arduino Duemilanove (Arduino, Turin, Italy) sent 3 Hz pulses to Power-LED-Module MinoStar (2.37 W, 36 lm, 30 °, Signal Construct GmbH, Niefern-Oeschelbronn, Germany), which produced light at an intensity of 1.64 mW/mm^2^ for 36 minutes. All constructs were prepared using either T4 DNA Ligation (New England Biolabs, Ipswich, MA, USA) or HiFi DNA assembly (New England Biolabs, Ipswich, MA, USA). Primer sequences for *inx-18int2* and *lim-6int3s* (for specific expression in AVG and in PVT, respectively; Chien et al., 2017; Oren-Suissa et al., 2016) were adjusted to fit our cloning strategy.

The following strains were generated and/or used:

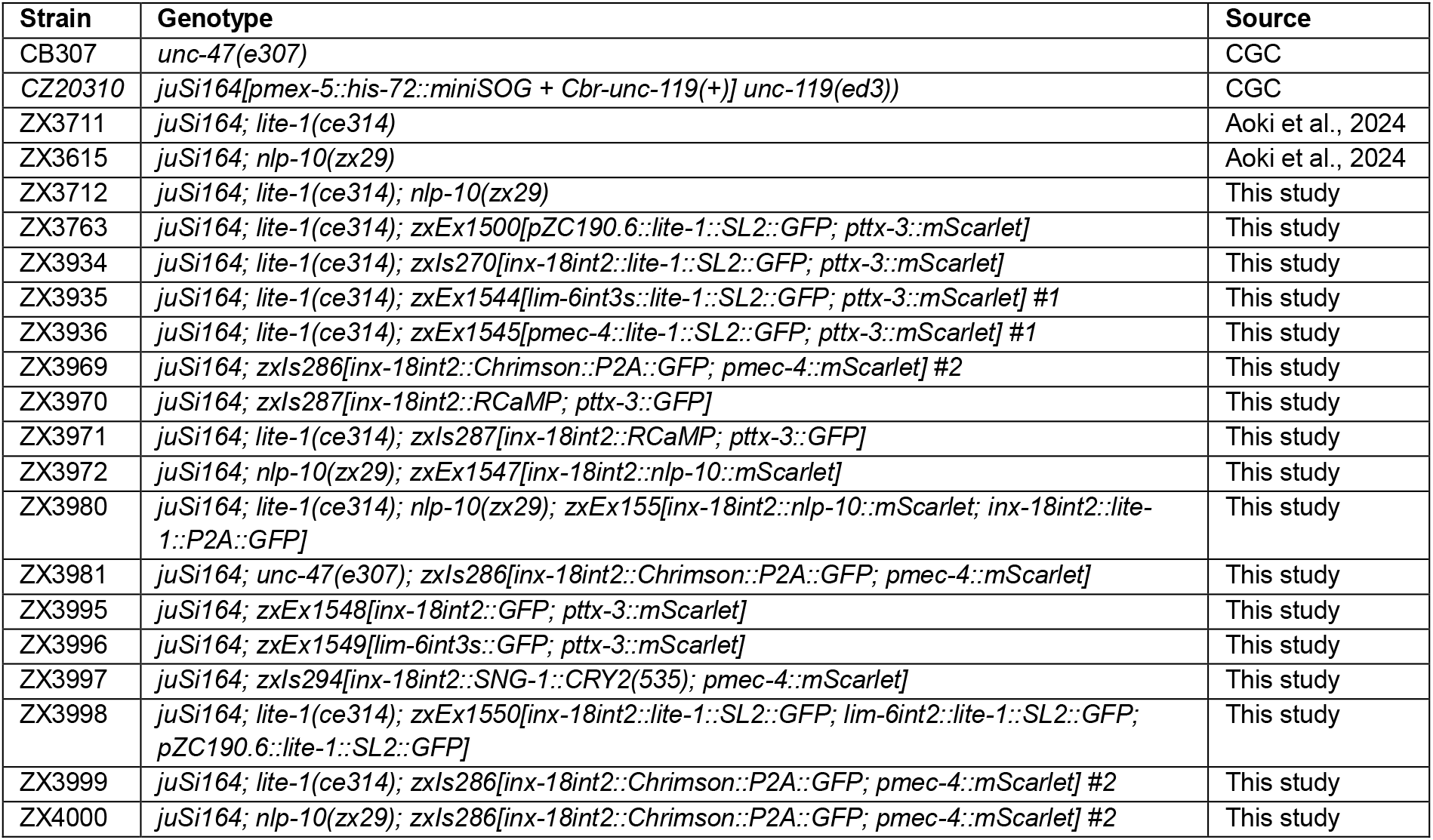

### Behavioral assays

20-40 L4 animals were picked on NGM plates 16 h prior to experiments (performed on the following day), unless stated otherwise. Plates were kept at 22 °C in the dark. Crawling behavior of animals was recorded with the Multi-Worm-Tracker (MWT), as previously described (Swierczek et al., 2011; Aoki et al., 2024). For the measurements investigating the response to blue light, an LED ring illuminated the plate with 1.0 mW/mm^2^ at 470 nm. Where appropriate, activation of Chrimson was performed with an intensity of 0.3 mW/mm^2^ at 620 nm wavelength. After the initial data processing by the Choreography software, which evaluates the speed and body bending of the animals, the data was analyzed and visualized with the help of custom MATLAB scripts (Aoki et al., 2024). Baseline speed was normalized to 30 s before (first) blue light illumination. For each measurement (indicated as N), the mean speed of recorded animals (indicated as a range for the number of animals n) was used for further analysis. All figures presenting crawling speed show mean values ± standard error (SEM). Box or violin plots represent the median of the means (N measurements) with 25 / 75 % quartiles. Means for each measurement were plotted as individual scatters.

Representative videos of strains were recorded with a LabVIEW-based custom software MS-Acqu under the same conditions as measurements were done. Displaying recording time and indication of the illumination period, as well as the compression of the original videos were performed with Clipchamp (Microsoft).

### Fluorescence imaging

Animals were placed on pads composed of 2.5 % agarose dissolved in M9 buffer and immobilized with Levamisole-hydrochloride (20 mM; Sigma-Aldrich, St. Louis, Missouri, USA) and 0.1 µm PolyBeads (Polysciences Inc., Warrington, Pennsylvania, USA). Images were obtained with a Kinetix22 camera (Teledyne Photometrics, Tucson, Arizona, USA), mounted on a Zeiss AxioObserver Z1 with a 40x oil immersion objective (EC Plan-NEOFLUAR 40x/1.3 Oil DIC. 420462-9900. Carl Zeiss, Oberkochen, Germany). Fluorescent proteins were excited with 460 or 590 nm Prior Lumen LEDs (Lumen 100, Prior Scientific, Cambridge, UK), which were regulated by µManager 2.0. An AOTFcontroller script (https://github.com/micro-manager/mmCoreAndDevices/blob/main/DeviceAdapters/Arduino/AOTFcontroller/AOTFcontroller.ino) was applied to control Z-stack recordings with an Arduino Uno.

To obtain representative images of fluorescent signal an EGFP/mCherry double band pass filter (450-490 nm and 555-590 nm excitation, 520/630 nm beamsplitter and 520/20; 630/30 nm emission; AHF Analysentechnik, Germany) was used. For the verification of LITE-1::SL2::GFP expression in ASK, animals were incubated in DiI staining solution (Invitrogen, Thermo Fisher Scientific Inc, Darmstadt, Germany). A 1 mM stock solution was diluted 1:100 in M9 buffer, and young adult animals were transferred into the staining solution and rotated for four hours in the dark. Before imaging, animals were washed twice in M9 to remove dye residues. Schematic representations of whole animals with the position of AVG, ASK or PVT were created in BioRender.

### Calcium imaging assays

Animals expressing AVG::jRCaMP1b were prepared in the same manner as for the fluorescence imaging and observed under a Zeiss AxioObserver Z1 with a 100x oil immersion objective (EC Plan-NEOFLUAR 100x/1.3 Oil DIC. 420490-9900), and equipped with an RCaMP filter set (479/585 excitation, 605 nm beamsplitter and 647/57 nm emission, AHF Analysentechnik, Germany). 200 ms light pulses (2 Hz) at 590 nm were used to excite the jRCaMP1b signal, while additional 200 ms light pulses (2 Hz) at 460 nm (0.34 mW/mm^2^) were applied between 30 and 40 seconds of recordings, to evoke the noxious light response. A custom script running on Arduino created an illumination pattern, to solely obtain images during yellow light illumination. For quantitative analysis, a ROI was placed along the axon of AVG, while a second ROI was placed on the body wall of animals, near the AVG axon. Signal intensity was measured with Fiji (ImageJ 1.53c, National Institutes of Health, USA). Afterwards, the intensity values obtained from the second ROI were subtracted from those of the first ROI, as background correction. Baseline values were calculated 10 s before the start of the illumination with blue light and the signal intensity is presented as follows:

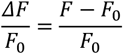

### Statistical analysis

Statistical analysis, as well as presentation of obtained data in graphs was done using Prism 9.4.1 (Graphpad software). For comparisons of more than two datasets, one- or two-way ANOVAs were applied with Tukey as post-hoc analysis. For comparisons of two normally distributed datasets, an unpaired t-test was applied. For comparison of datasets that were not normally distributed, a Mann-Whitney test was applied (Fig. 3F). Asterisks indicate the significance levels of p < 0.05 (*), p < 0.01 (**), p < 0.001 (***) and p < 0.0001 (****).

## Data Availability Statement

Data is mostly on-line measurements from multiworm tracker, for which no video is saved. These data are summarized in Supplementary Data File 1, sorted for each figure of the paper. All imaging data is available from the authors and will be made publicly available in an online repository, e.g. FigShare. Strains and plasmids are available upon request. Matlab scripts used to analyze multiworm tracker data can be found at Zenodo (https://doi.org/10.5281/zenodo.13958554).

## Author contributions

E.D., I.A. and A.G. planned experiments. E.D. designed and performed experiments, wrote and edited the manuscript. I.A. and A.G. supervised the work. I.A. and A.G. edited the manuscript. A.G. finalized the manuscript for publication, and secured funding.

## Acknowledgements

We are grateful for expert technical assistance to Katharina Kuhlmeier, and for help with writing code and data analysis to Dennis Rentsch and Noah Schuh. Some strains were obtained from the *Caenorhabditis* Genetics Center, which is funded by the NIH Office of Research Infrastructure Programs (P40 OD010440).

## Funding

This work was funded by the Deutsche Forschungsgemeinschaft (DFG), grant GO1011/18-1, and by core funding from Goethe University, to A.G.

## Conflict of interest

The authors declare no competing interests.

